# Massively parallel sequencing of 165 ancestry-informative SNPs and forensic biogeographical ancestry inference in three southern Chinese Sinitic/Tai-Kadai populations

**DOI:** 10.1101/2020.12.05.412866

**Authors:** Guanglin He, Jing Liu, Mengge Wang, Xing Zou, Tianyue Ming, Sumin Zhu, Hui-Yuan Yeh, Chuanchao Wang, Zheng Wang, Yiping Hou

**Author notes:** Corresponding Author: Zheng Wang and Yiping Hou, **Zheng Wang**, Institute of Forensic Medicine, West China School of Basic Medical Sciences and Forensic Medicine, Sichuan University, Chengdu 610041, China, Address: 3-16 People’s South Road, Chengdu 610041, China, **Yiping Hou**, Institute of Forensic Medicine, West China School of Basic Medical Sciences and Forensic Medicine, Sichuan University, Chengdu 610041, China, Address: 3-16 People’s South Road, Chengdu 610041, China. The author contributed equally to this work and should be considered co-first author (Guanglin He, Jing Liu).

## Abstract

Ancestry informative markers (AIMs), which are distributed throughout the human genome, harbor significant allele frequency differences among diverse ethnic groups. The use of sets of AIMs to reconstruct population history and genetic relationships is attracting interest in the forensic community, because biogeographic ancestry information for a casework sample can potentially be predicted and used to guide the investigative process. However, subpopulation ancestry inference within East Asia remains in its infancy due to a lack of population reference data collection and incomplete validation work on newly developed or commercial AIM sets. In the present study, 316 Chinese persons, including 85 Sinitic-speaking Haikou Han, 120 Qiongzhong Hlai and 111 Daozhen Gelao individuals belonging to Tai-Kadai-speaking populations, were analyzed using the Precision ID Ancestry Panel (165 AISNPs). Combined with our previous 165-AISNP data (375 individuals from 6 populations), the 1000 Genomes Project and forensic literature, comprehensive population genetic comparisons and ancestry inference were further performed via ADMIXTURE, TreeMix, PCA, *f*-statistics and N-J tree. Although several nonpolymorphic loci were identified in the three southern Chinese populations, the forensic parameters of this ancestry inference panel were better than those for the 23 STR-based Huaxia Platinum System, which is suitable for use as a robust tool in forensic individual identification and parentage testing. The results based on the ancestry assignment and admixture proportion evaluation revealed that this panel could be used successfully to assign individuals at a continental scale but also possessed obvious limitations in discriminatory power in intercontinental individuals, especially for European-Asian admixed Uyghurs or in populations lacking reference databases. Population genetic analyses further revealed five continental population clusters and three East Asian-focused population subgroups, which is consistent with linguistic affiliations. Ancestry composition and multiple phylogenetic analysis further demonstrated that the geographically isolated Qiongzhong Hlai harbored a close phylogenetic relationship with Austronesian speakers and possessed a homogenous Tai-Kadai-dominant ancestry, which could be used as the ancestral source proxy in population history reconstruction of Tai-Kadai-speaking populations and as one of the representatives for forensic database establishment. In summary, more population-specific AIM sets focused on East Asian subpopulations, comprehensive algorithms and high-coverage population reference data should be developed and validated in the next step.

## 1. Introduction

Genetic markers with significant allele frequency differences among geographically/ethnically diverse populations have been widely utilized to explore population stratification and dissect population movement and admixture in anthropology, archeology and population genetics [1, 2]. These genetic markers used for the identification of genetic stratification in case-control genome-wide association studies and for biogeographical ancestry inference in forensic science are referred to as ancestry-informative markers (AIMs). In recent years, many AISNP sets have been developed to infer the biogeographic ancestry of unknown individuals by the forensic community [1, 3-8]. However, AISNP sets differ in suitability for widespread application due to their initial development purpose focused on specific targeted populations and the lack of global high-coverage reference population data [9, 10]. Recently, Thermo Fisher Scientific released a commercial ancestry panel, the Precision ID Ancestry Panel, based on the massively parallel sequencing (MPS) platform. This MPS panel includes 165 AISNPs; 55 of these markers were selected from Kidd’s AIM set, and 123 markers were selected based on Seldin’s AIM set [5, 11, 12]. In the past few years, forensic efficiency parameters of this AISNP panel have been validated in multiple populations, and their application for ancestry inference has been investigated in Europeans, Africans, Americans and a subset of northwestern East Asians [3, 4, 6, 12-15].

China, with a population size of over 1.4 billion, is enriched in ethnic, cultural, linguistic and genetic diversity. There are at least nine language families, including the world’s second-largest, the Sino-Tibetan (Tibetan-Burman and Sinitic) [16], as well as the Austronesian, Austroasiatic, Hmong-Mien, Tai-Kadai and Trans-Eurasians (Mongolic, Tungusic and Turkic) families and others. In addition, East Asia has two independent animal/plant domestication centers, with the northern center localized in the upper and middle Yellow River Basin where foxtail and broomcorn millet were domesticated and the southern center localized in the Yangtze River Basin where wet rice (*Oryza japonica*) was domesticated [17]. Complex population history, including special patterns of Pleistocene-Holocene transitions, processes of neolithization and agriculture-mediated Holocene population expansion, has shaped the modern genetic diversity of East Asia [18, 19]. Previous genetic studies have demonstrated a significant north-south genetic distinction in China [19] and significant genetic differences between Tibeto-Burman-, Turkic-, Sinitic- and Tai-Kadai-speaking populations [20-23]. Recent paleogenetic evidence has also suggested that significant genetic differentiation between northern and southern East Asians and between Highlanders and lowland East Asians occurred from the early Neolithic period [22, 24]. This differentiated demographic history of geographically separated East Asians and their specific genetic structures also suggests the potential for subpopulation identification in forensic ancestry inference. Thus, East Asian-specific AIM panels and corresponding population reference databases need to be constructed for forensic practice. Our previous studies were focused on the exploration of population reference data and evaluation of ancestry inference of Tibeto-Burman- and Turkic-speaking populations using the Precision ID Ancestry Panel [4, 15]. Ancestry inference efficiency in southern Chinese populations has remained in its infancy. This region is not only the original center of the Austronesian-, Austroasiatic-, Tai-Kadai- and Hmong-Mien-speaking populations but also the massive genetic admixture cradle between the southward spreading Han Chinese and indigenous populations[25]. Thus, this study focused on three main ethnic groups from southern China, including the Han and Hlai from Hainan Island and the Gelao from Guizhou Province. Hainan Island, separated from the mainland by the Qiongzhou Strait, is the southernmost and smallest province in China. The Han and Hlai ethnic groups account for over 98% of the population on the island. The Han Chinese, the largest ethnic group in China, is mainly concentrated in the northeast, north and coastal areas of Hainan Island, and many of their ancestors migrated from the mainland during the Song dynasty. The Hlai people are an indigenous community on the island with considerable differences in language, culture and origin compared with the Han people. The Hlai people originally had their own specific language, which has been regarded as a primary branch of the Tai-Kadai language family [25, 26]. Another Tai-Kadai-speaking population, the Gelao, mainly (over 96%), reside in Guizhou Province.

Here, to provide a better understanding of the genetic backgrounds of the Tai-Kadai-speaking Hlai and Gelao ethnic groups and southern Han Chinese and to verify the feasibility of the Precision ID Ancestry Panel for inferring biogeographic ancestors of southern East Asians, we generated population data for 165 AISNPs among 316 southern Chinese individuals from Tai-Kadai-speaking Hlai and Gelao and Sinitic-speaking Han populations using the Ion S5 XL system and co-analyzed these data with different reference datasets in three sets of comprehensive population structure analyses. First, we explored the forensic characteristics of 165 AISNPs via the STRAF and HID SNP Genotyper. Second, to explore the genetic affinity between southern Chinese populations and worldwide reference groups, we merged newly generated data of 164 AISNPs (except for rs10954737, which is lacking in the 1000 Genomes Project data) with six populations from our previous studies [4, 27], Kazakhs [12], 26 populations from the 1000 Genomes Project [28] and 76 populations from the 170-AISNPs-Kidd-Seldin dataset (**Table S1** and **Fig. 1A**) [29], which is the highest coverage dataset and consists of 6,933 individuals from 112 worldwide populations. Comprehensive population genetic analyses were conducted based on genotype-based analysis via principal component analyses (PCA), model-based ADMIXTURE, F_st_, distance-based TreeMix modeling, admixture-*f*_*3*_ statistics, outgroup-*f*_*3*_ statistics, four population-based symmetrical *f*_*4*_-statistics and neighbor-joining (N-J) phylogenetic trees. Third, to further explore genetic differences and similarities between the newly generated East Asians and additional reference populations focused on two AISNP subsets, we performed two sets of frequency-based analyses: one using the Kidd-AIM set with 157 populations based on 55 AISNPs and one using the Seldin-AIM set with 140 populations based on 123 AISNPs. Genetic affinity was estimated via PCA, pairwise genetic distance and the N-J tree. Our research here mainly focused on the evaluation of the effectiveness of the 165-AISNP panel, providing an additional population-specific database, exploring the population substructures within East Asians and reconstructing their potential genetic admixture and gene flow events.

**Fig. 1.**
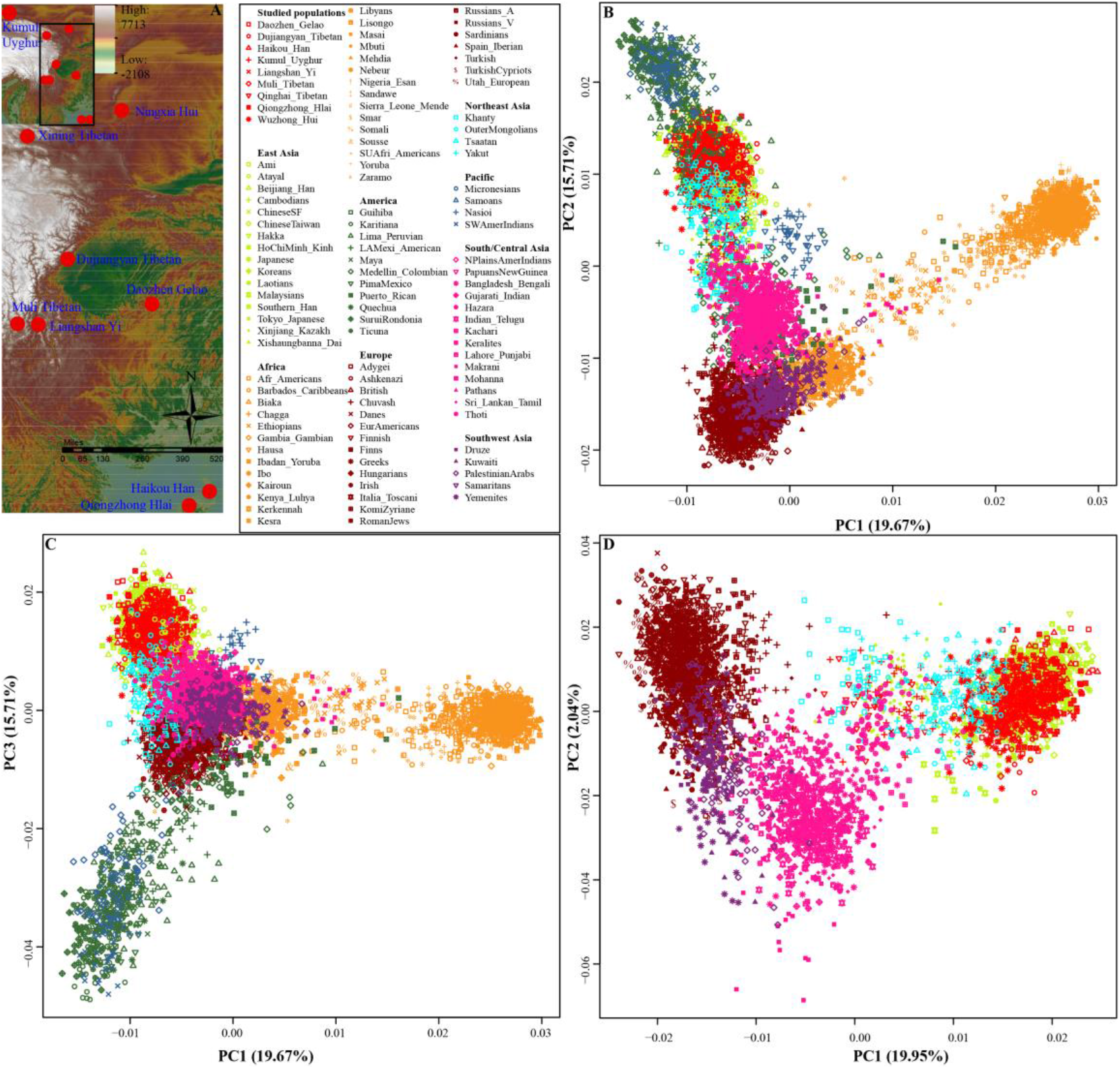
Overview of genetic affinity. (**A**). Geographical positions of seven newly generated and previously investigated populations. (**B∼C**). Principal component analysis (PCA) among 6933 individuals from 112 populations based on the top two components and the combination of the first and second components. (**D**). Two-dimensional PCA plots among 4506 individuals from 67 populations reconstructed based on the first two components. All included individuals were color-coded by geographical divisions.

## 2. Materials and methods

### 2.1. Sample preparation

Human blood samples were collected with the approval of the Ethics Committee at Sichuan University (Approval Number: K2019040). Peripheral blood was collected from each donor after receiving written informed consent. This study followed the ethical principles stated in the Helsinki Declaration of the World Medical Association. Blood samples were collected from 316 Chinese persons, including 85 Haikou Han, 120 Qiongzhong Hlai and 111 Daozhen Gelao individuals. A geographical map of the three ethnic groups is shown in **Fig. S1**. All included participants were self-declared indigenous members of the corresponding ethnicity groups or had lived in sample collection locations for at least three generations.

Human genomic DNA was extracted using the PureLink Genomic DNA Mini Kit (Thermo Fisher Scientific, USA) or QIAamp DNA Blood Mini Kit (QIAGEN, Germany) and quantified using the Quantifiler Human DNA Quantification kit (Thermo Fisher Scientific) on an Applied Biosystems 7500 Real-time PCR System (Thermo Fisher Scientific) following the manufacturer’s instructions. DNA samples were normalized to 1.0 ng/μL and stored at -20 °C until library amplification.

### 2.2. Library construction, template preparation and sequencing

AISNP libraries were prepared according to the Precision ID Ancestry Panel protocol (Revision C), as described in detail in our previous studies [4, 15, 21, 26]. Briefly, 1.0 ng of template DNA was amplified, and then the amplicons were partially digested using FuPa reagent. Next, barcode adapters were ligated to the amplicons, and the resulting libraries were purified with Agencourt AMPure XP Reagent (Beckman Coulter, USA). Diluted libraries (1:100 dilution) were quantified with the Ion Library TaqMan Quantitation Kit (Thermo Fisher Scientific), and the normalized libraries were pooled for template preparation on the Ion Chef Instrument. The sequencing reaction was performed on the Ion S5 XL System according to the manufacturer’s protocol (https://assets.thermofisher.com/TFS-Assets/LSG/manuals/MAN0017767_PrecisionID_SNP_Panels_S5_UG.pdf).

### 2.3. Sequencing data acquisition and analysis

Preliminary analysis of all sequencing data was automatically conducted using Torrent Suite software V5.10.0 (Thermo Fisher Scientific) with the *Homo sapiens* hg 19 genome as the alignment reference. The HID SNP Genotyper Plugin was used for secondary analysis with the target region file (PrecisionID_AncestryPanel_targets.bed) and the hotspot region file (PrecisionID_AncestryPanel_hotspots.bed), using the default settings.

Allele frequencies of 165 AISNPs and corresponding forensic statistical parameters, including genetic diversity (GD), observed heterozygosity (Ho), match probability (MP), power of discrimination (PD), polymorphism information content (PIC), power of exclusion (PE) and typical paternity index (TPI), were calculated using the online tool STR Analysis for Forensics (STRAF) [30]. Arlequin was used to calculate the p-values for Hardy-Weinberg equilibrium and linkage disequilibrium [31].

### 2.4 Data merging and statistical analysis

Three different datasets were used for population genetic analyses: one raw genotyped dataset and two allele-frequency-based datasets (the Kidd and Seldin panels). Initially, in this study we merged 316 genotypes generated from western Chinese Tibeto-Burman-speaking Tibetan and Yi, northwestern Turkic-speaking Uyghur and Sinitic-speaking Hui populations from our previous studies [4, 15] with Kazakh population data provided by the Zhu Lab [12] and 76 populations from the 170-AISNPs-Kidd-Seldin dataset [29]. Then, we combined the above data with 26 populations worldwide from the 1000 Genomes Project [28] via the mergeit package. Detailed information about the studied populations is shown in **Table S1**. Furthermore, allele frequency data for the Kidd and Seldin SNPs [4, 5, 10, 15, 32] were merged to investigate population similarities and differences. The Kidd dataset (55 AISNPs) includes 157 populations of Africans, Central and South Asians, East Asians, Europeans, North Americans, Oceanians and South Americans. Seldin’s dataset includes 140 populations of Africans, Central and South Asians, East Asians, Europeans, North Americans, Oceanians and South Americans [33] (**Table S1**).

The pairwise Cavalli-Sforza genetic distances were calculated using PHYLIP software, and the fixation index (pairwise F_st_ genetic distance) was evaluated using STRAF [34]. Principal component analysis (PCA) was performed based on genotypic data using Plink and SmartPCA [35, 36]. PCA based on allele frequency distributions was conducted using MVSP software. Model-based ADMIXTURE was used to infer individual- and population-level ancestry composition, and the additional parameters were set to - B100 and --cv = 10 [37, 38]. The phylogenetic relationship tree was reconstructed using Mega7 [37, 38]. Multidimensional scaling analysis was conducted using R. Admixture-*f*_*3*_-statistics in the form *f*_*3*_*(Source1, Source2; studied populations)* were analyzed to explore the admixture signals, and outgroup-*f*_*3*_-statistics in the form *f*_*3*_*(Reference population1, reference population2; Mbuti)* were applied to explore the shared genetic drift using the *qp3Pop* program in ADMIXTOOLS [38]. Symmetrical *f*_*4*_-statistics in the form *f*_*4*_ *(Reference population1, reference population2; studied population, Mbuti)* and *f*_*4*_*(Reference population1, studied population; reference population2, Mbuti)* were estimated using D-statistics [38] with the additional parameter of the *f*_*4*_ model: YES.

### 2.5. Quality control

Control DNA 007 (Thermo Fisher Scientific) and RT-PCR Grade Water (Thermo Fisher Scientific) were used as the positive and negative controls, respectively, for each batch of library construction, template preparation and sequencing. All experiments were conducted at the Forensic Genetics Laboratory of the Institute of Forensic Medicine, Sichuan University, which is an accredited laboratory (ISO 17025), in accordance with quality control measures. Additionally, the laboratory has been accredited by the China National Accreditation Service for Conformity Assessment (CNAS). All genotypes generated in this study were confirmed by two scientists. We strictly followed the recommendations of the Chinese National Standards and Scientific Working Group on DNA Analysis Methods [39].

## 3 Results

### 3.1 Forensic relative statistical parameters

All sequencing data generated in this study were analyzed using Torrent Suite Software, and the HID SNP Genotyper Plugin was used for secondary analysis with the default analysis settings. After removing the samples with genotypes that did not fulfill the quality thresholds (denoted ‘NN’) and “no calls” (denoted ‘NOC’) as determined by the plugin, a total of 316 individuals were included in the subsequent analyses. The detailed genotypes of 165 AISNPs from 316 individuals are listed in **Table S2**. For the three studied populations, no significant deviation from Hardy-Weinberg equilibrium was found at 165 sites after Bonferroni correction (p > 3.0303 × 10^−4^), and no significant deviation from linkage disequilibrium was identified among all pairwise sites after correction for multiple tests (p > 3.70 × 10^−6^). As expected, not all 165 AISNPs were highly polymorphic in Chinese populations; 12, 10 and 15 SNPs were monomorphic in Daozhen Gelao, Haikou Han and Qiongzhong Hlai, respectively (marked with bold font in **Tables S3-S5**). The combined match probability (CMP) values in Haikou Han, Qiongzhong Hlai and Daozhen Gelao were 1.46E-46, 5.54E-46 and 2.68E-47, respectively, and the combined power of exclusion (CPE) values were 0.999999978, 0.999999968 and 0.999999965, respectively. These observed combined values of discriminatory power were greater than the combined forensic features of the Huaxia Platinum commercial kit [18, 40], which indicates that this ancestry-inference panel could be used for forensic personal identification.

### 3.2 Forensic ancestry inference via HID SNP Genotyper

Admixture prediction and population likelihoods were calculated via the ancestry prediction plugin built into Torrent Suite software. All individuals included were able to be successfully assigned to East Asian origin. However, when we focused on subpopulation origins, misassignment occurred due to the lack of corresponding reference data or discriminatory resolution limitations of these AIMsets. **Fig. S2** presents an example to demonstrate that continental discrimination could be obtained using this ancestry panel, but there were limitations in distinguishing intracontinental subpopulations. A Haikou Han was assigned to the admixture of 90% East Asian-related ancestry, 5% South Asian-related ancestry and 5% American-related ancestry. The predicted log-likelihood ratio value reached 106.31. Focusing on the subpopulation origin, the prediction result showed that this individual was most likely to have originated from the Taiwanese Han, with a likelihood ratio of 2.26E-46, followed by the Hakka, HapMap Han, Japanese and Korean subpopulations.

### 3.3 Population structure and genetic background revealed by raw genotype data

To further explore the ancestry discrimination power of this ancestry panel, we merged our data [4, 27] with data from 26 worldwide reference populations in the 1000 Genomes Project [28], 76 populations from the 170-AISNPs Kidd-Seldin dataset [29] and one Xinjiang Kazakh [12]. We performed a principal component analysis of 6,933 individuals from 115 worldwide populations. As shown in **Fig. 1B**, a two-dimensional plot based on PC1 and PC2 was able to assign all included populations into two genetic clines (African cline and Eurasian cline). The African cline began with southern Africans on one side and ended with Europeans at the other end, and the Eurasian cline started from Native Americans and ended with Europeans. East Asians overlapped with our studied populations and clustered with each other at a position between the Native American cluster and the South/Central Asian cluster. PC3 accounted for 15.71% of the variance and could with statistical significance separate East Asians and Native Americans from others (**Fig. 1C**). Focusing on 69 Eurasian populations (**Fig. 1D**), we identified three clusters. Europeans were clustered together, located in the upper left of the two-dimensional scatterplot, while South Asians were clustered in the middle-lower position and East Asians in the right middle position of the scatterplot. PC1 revealed the genetic differences between East Asians and other worldwide groups. Focusing only on the reference populations from the 1000 Genomes Project, we found four clearly separated population clusters, which indicated that biogeographical ancestry inference within intercontinental contact regions or mixed populations is another difficult problem (**Figs. S3-4**).

We calculated the pairwise F_st_ genetic distances among 112 populations. As shown in **Figs. S5-8** and **Table S6**, Haikou Han shared a close genetic relationship with Hakka (0.0009), followed by Southern Han (0.0011), Daozhen Gelao (0.0031) and Ho Chi Minh Kinh (0.0031). Genetic distances between Haikou Han and other East Asians were also relatively small, all less than 0.0996. The F_st_ genetic distance between Qiongzhong Hlai and Xishuangbanna Dai was the smallest (0.0051), followed by distances from Ho Chi Minh Kinh (0.0060), Laotians (0.0086), Haikou Han (0.0092) and Southern Han (0.0137). Daozhen Gelao shared a close genetic relationship with Hakka (0.0001) and Beijing Han (0.0005), followed by Southern Han (0.0014). The overall genetic distances were smallest between European-Asian admixed populations and other included worldwide reference populations, such as Xinjiang Kazakhs (0.1545 ± 0.1181) and Kumul Uyghurs (0.1632 ± 0.1256). Based on the heatmap results, clustering patterns of genetic differences and similarities were not only in accordance with continental divisions but also consistent with linguistic affiliations. As shown in **Fig. S6**, Austronesian-speaking Ami and Atayal clustered, and Tibeto-Burman-speaking Tibetan and Yi populations also clustered first. The identified population genetic affinities were confirmed via the N-J-based phylogenetic clustering results (**Figs. S9∼10**).

The ancestry coefficient was first estimated via model-based ADMIXTURE from 6,399 individuals worldwide. Cross-validation error results suggested that the five-source model with the least error (0.5146) could be used to explain the continental genetic variations. South Asian ancestry was maximized in Telugu Indians (0.783), East Asian ancestry was maximized in Austronesian Ami, African ancestry was maximized in Ibadan Yoruba (0.99), Native American ancestry was maximized in Karitiana (0.978) and European ancestry was maximized in the Irish (0.865). As shown in **Fig. 2A, Fig. S11** and **Table S7**, all East Asians were modeled as deriving principal ancestry from the Ami-related ancestry proxy with additional admixture from neighboring continental sources. Detailed ancestry admixture proportions at the individual and population levels were further evaluated among the newly generated East Asians and twenty-six 1000-genome-project populations with 2-7 predefined ancestral populations (**Fig. S7**). When we assumed two ancestral populations (K = 2), the blue ancestral component and the orange ancestral component were identified. The blue ancestral component had the highest proportion in the Eurasian group, and the percentage exceeded 99% in three East Asian groups and one European group. The orange ancestral component appeared in the highest proportion in the African population and could be regarded as an African-specific ancestral component. When three ancestral populations were assumed, the western Eurasian ancestral population was independently modeled and separated from other Eurasians. The yellow component could be regarded as the European-specific ancestral component. The orange ancestral component remained the African-specific ancestral component, while the blue component appeared maximally in East Asians. South Asians and Americans were modeled as a mixed population of two or three ancestral components. It is worth noting that the Kumul Uyghur, Xining Tibetan and Xinjiang Kazakh subpopulations were simulated as a mixed population of eastern and western Eurasian peoples. Under the model of four ancestral groups, we were able to identify a new South Asian-specific ancestral component (the pink component), which appeared in the highest proportion in the Indian Telugu subpopulation (0.82). Orange African-specific ancestry was maximized in Ibadan Yoruba (0.99). Yellow European-specific ancestry was maximized in British individuals (0.87), and blue East Asian-specific ancestry was maximized in Beijing Han individuals (0.93). Cross-validation errors revealed that the proximal K value was 5. Five large intercontinental ancestral groups were simulated: African, European, South Asian, East Asian and American. The green ancestral component appeared here for the first time and was maximized in Lima Peruvians (0.69), followed by Los Angeles Mexican Americans (0.42), Medellin Colombians (0.23) and Puerto Ricans (0.13). When six ancestral populations were assumed, the Tibetan-specific ancestral component was gradually separated from the others. As the K values further increased, additional substructures within intercontinental groups were gradually molded.

**Fig. 2.**
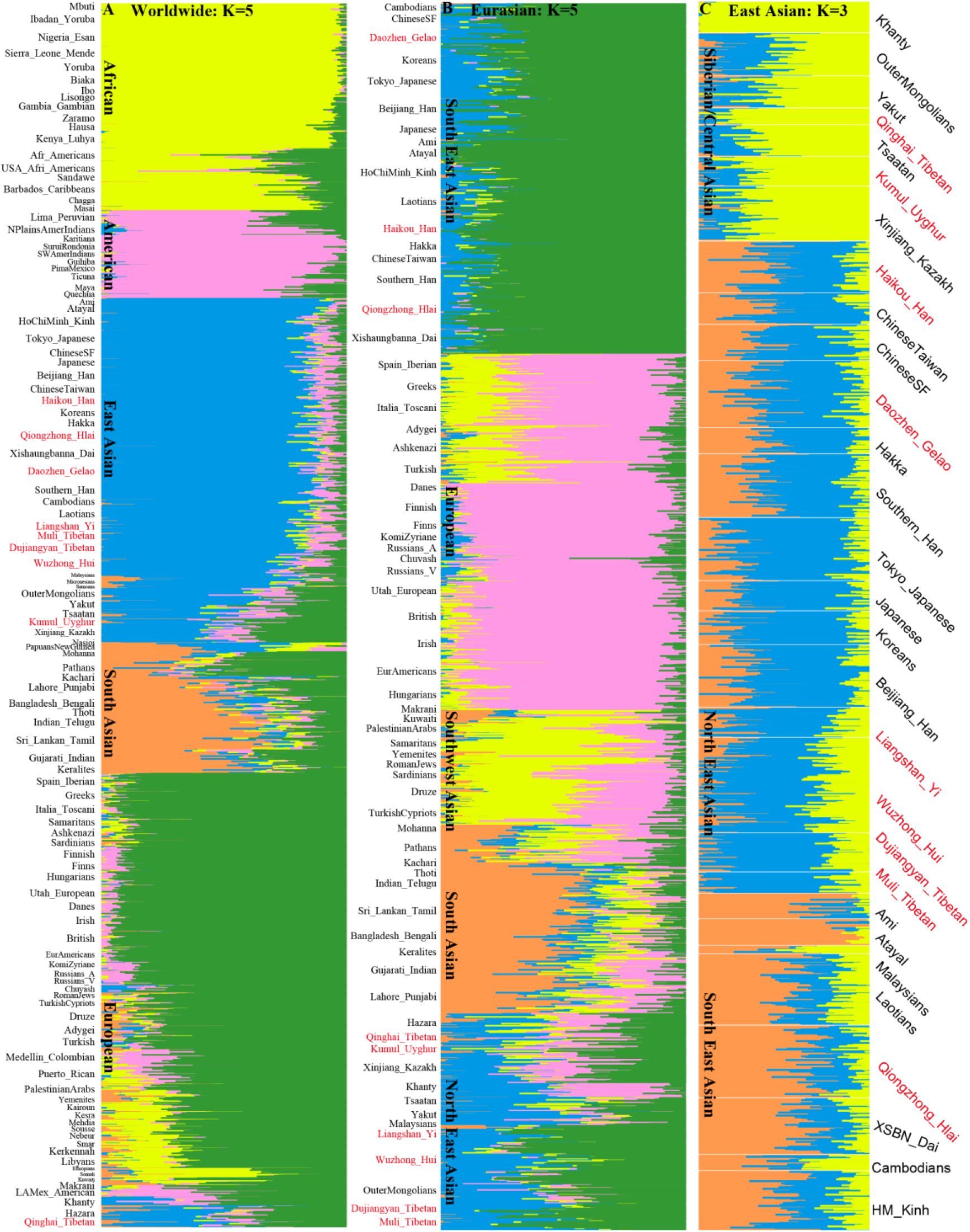
Model-based ADMIXTURE results displaying ancestry composition. (**A**). Ancestry composition among 112 populations worldwide with five predefined source populations. (**B**). Ancestry coefficient among 506 individuals from 67 populations with the least error (K = 5). (**C**). Individual bar plots from ADMIXTURE analyses at K = 3 showed three substructures among East Asians.

We also dissected the ancestry components within the region-specific population subsets (**Fig. 2B∼C, Figs. S13∼14 and Tables S8∼9**). 4,506 Eurasian individuals from 68 populations could be clustered into five subgroups: a Southern Asian subgroup represented by Telugu Indians (0.688), a northern East Asian subgroup represented by Yakut (0.639) and other Tibeto-Burman speakers, a southwestern Asian subgroup represented by Samaritans (0.759), a European subgroup represented by Finns (0.835) and a southern East Asian subgroup represented by Austronesian speakers (0.873 for Atayal and 0.817 for Ami) and Tai-Kadai speakers (0.771 for Hlai and 0.739 for Dai). Here, we found that three studied populations from southern China were primarily formed by indigenous southern ancestral sources and then obtained additional admixtures from northern East Asians. Northern East Asia harbored greater Yakut-related or Tibetan-related ancestry and less southern Tai-Kadai-related or Austronesian-related ancestry. Further substructures within East Asians could also be identified via ADMIXTURE among 30 eastern Eurasian populations (**Fig. 2C**). Populations from southern China, northern China and northwestern China possessed their region/ethnicity-specific ancestries, and these genetic signatures also reflected their differentiated demographic histories [19, 22, 23].

### 3.4 Population splits and gene flow revealed by TreeMix Analysis

To further explore the evolutionary relationships from the perspectives of worldwide reference populations, we performed distance-based TreeMix analysis between the newly generated East Asians and 26 other previously published 1000 Genomes Project groups. We constructed the maximum likelihood tree with Ibadan Yoruba as the root group, and 0 to 16 mixture events were gradually simulated to reconstruct the population splits and gene flow events. As shown in **Fig. S15**, seven African populations were clustered together as the root, and five Europeans and five South Asians clustered as the western Eurasian cluster. Ten populations genotyped using the ancestry inference panel and five 1000-Genomes-Project populations clustered and formed the Eurasian branch. Lima Peruvians and Mexican Americans in Los Angeles gathered with Uyghurs, Kazakhs and other groups, which were located between Europeans and East Asians. Medellin Colombians and Puerto Ricans were clustered between the South Asian branch and the European Asian branch. Qiongzhong Hlai and Haikou Han in southern China first clustered with Beijing Han, Southern Han and Daozhen Gelao, successively and finally clustered with Tokyo Japanese, Xishuangbanna Dai and Ho Chi Minh Kinh. The above cluster formed the East Asian terminal cluster in the maximum likelihood tree. Dujiangyan Tibetan, Muli Tibetan and Liangshan Yi subpopulations clustered and formed the Tibetan-Burmese cluster. However, the Xining Tibetans and the Kumul Uyghur, Xinjiang Kazakh, and Ningxia Hui subpopulations formed a near-central Asian subcluster. It is worth noting that although Qiongzhong Hlai, Daozhen Gelao and Xishuangbanna Dai peoples belonged to the Tai-Kadai language family, they did not initially group in this maximum likelihood tree but first clustered with their geographically close populations.

When considering the potential gene flow influx among the included populations, as shown in **Fig. S15**, Kumul Uyghurs could be modeled as an admixture of 25.18 ± 1.82% Finn-related ancestry and 74.88 ± 1.82% East Asian-related ancestry (p < 2.225e-308), and Xinjiang Kazakhs were modeled as a mixture of 29.00 ± 3.15% European-related ancestry and 71.00 ± 3.15% Eastern Asian-related ancestry with the corresponding p-values of 0.0315. In the three genetic flow models (**Fig. S16**), Kumul Uyghur and Xinjiang Kazakh populations were both modeled as mixtures of Finn-related ancestral components (approximately 74%) and East Asian ancestral components (approximately 26%). Medellin Colombians were modeled as a mixture of 46.23 ± 2.95% Mexican-American-related ancestral components and 53.77 ± 2.95% European-related ancestral components with p-values less than 2.225e-308. When fifteen gene flow events were assumed, as shown in **Fig. S17**, 14.35 ± 1.10% non-African ancestral component, 17.74 ± 1.36% Lima Peruvian-related ancestral component and approximately 67.91% European-related ancestral component were mixed to form the Puerto Ricans with a p-value of less than 2.225e-308. The Kumul Uyghur people was modeled as a mixture of 40.11 ± 2.06% European ancestral component and 59.89 ± 2.06% East Asian ancestral components with p-values less than 2.225e-308. The Xinjiang Kazakhs were modeled as a mixture of 36.51 ± 2.47% European-related ancestral components and 63.49 ± 2.47% East Asian-related ancestral components. Xining Tibetans were modeled as a mixture of 51.08 ± 0.29% European-related ancestral components and 48.92 ± 0.29% East Asian-related ancestral components. The Sri Lankan Tamil population showed 26.24 ± 3.71% of its ancestral component associated with Utah Europeans. A total of 33.37 ± 3.00% of the European ancestral component was incorporated into the common ancestral population of the Medellin Colombians and the Mexican Americans in Los Angeles (p < 2.225e-308). A total of 43.01 ± 3.93% of the European ancestral component was pooled into the Colombian population in Medellin (p < 2.225e-308). A total of 14.74 ± 0.88% of the Pakistani-related ancestral component was mixed into the Barbados Caribbean (p < 2.225e-308). A total of 26.12 ± 1.04% of the Pakistani-related ancestral component was mixed into African Americans with a p-value less than 2.225e-308.

Focusing on Eurasian populations (**Fig. 3A**), patterns of population clustering were consistent with the ADMIXTURE modeling result. Western and eastern Eurasians were clustered at two ends of the tree, and populations from southern/central Asia were grouped and localized between them. Similar gene flow events from western Eurasians into northwestern East Asians were identified. Focusing on the clustering pattern within East Asians (**Fig. 3B**), we found that strong genetic affinity was associated with geographical division and linguistic affiliation. The overall patterns were consistent with the East Asian-based ADMIXTURE three-source model. Austronesian speakers and Tai-Kadai speakers were first grouped and then clustered with other southern East Asians, which formed the southern East Asian lineage. Tibetan-Burman speakers clustered and maintained a close genetic relationship with northern Chinese populations, which formed the northern East Asian lineage. Populations from northern Asia or northeast East Asia formed one clade with several different additional gene flow events. We finally used formal tests of the *f*-statistics to explore the shared ancestry or drift between three southern East Asian and other worldwide reference populations (**Tables S11∼13**). Statistically significant admixture-*f*_*3*_-statistics results focused on Daozhen Gelao and Haikou Han were mainly obtained when one source was from northern East Asians and the other was from southern East Asians. However, these signals were lacking when we used Qiongzhong Hali as the targeted population. These observed northern and southern genetic affinities were further confirmed via the obvious derived alleles shared between the studied populations and reference populations in *f*_*4*_ statistics (**Tables S14∼16**).

**Fig. 3.**
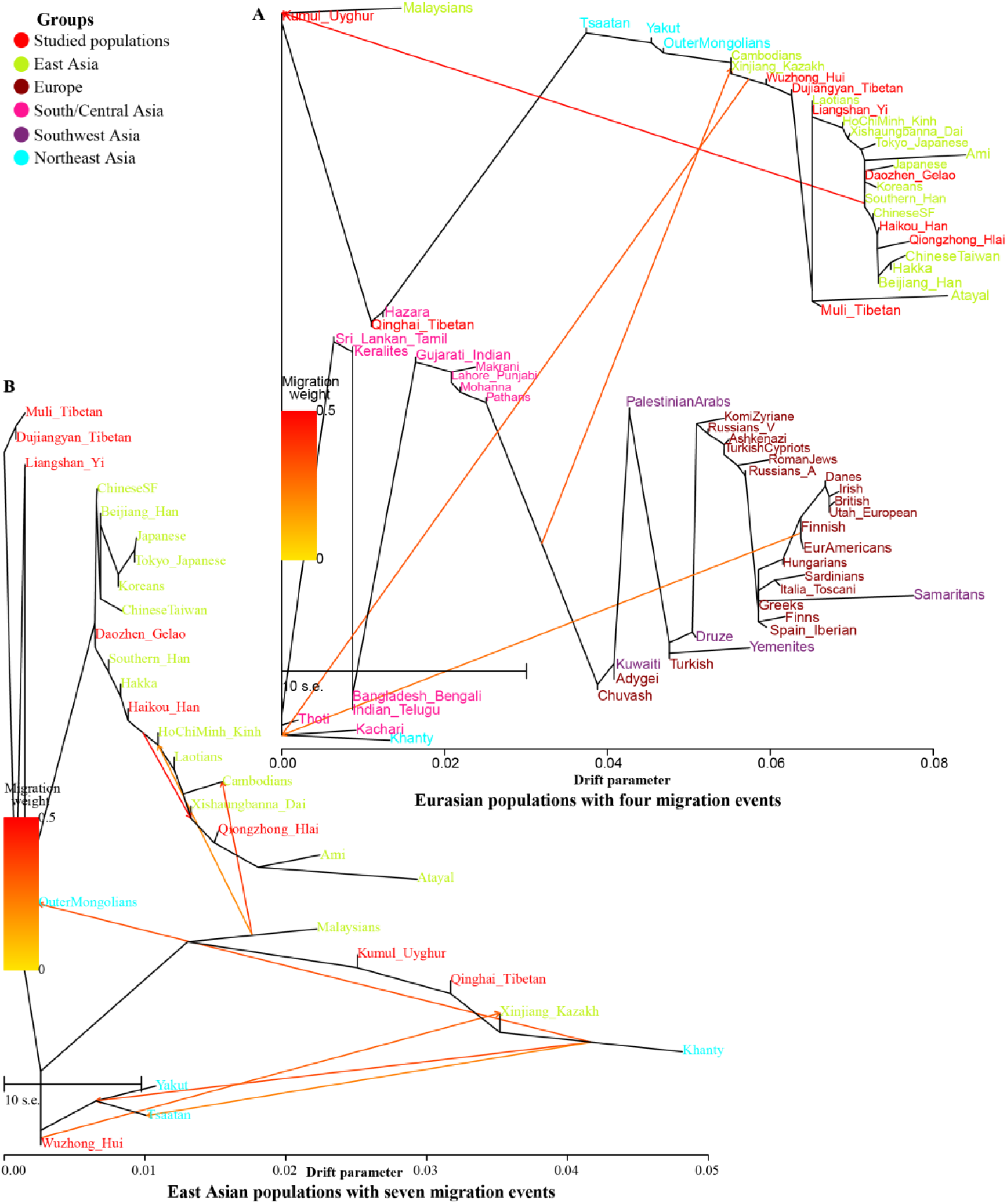
Inference of population separation and gene flow events. (**A**). Phylogenetic relationships and population splits were reconstructed by TreeMix under an assumption that four recent migrations among 67 Eurasian populations occurred. (**B**). East Asian-focused phylogenetic tree with seven gene flow events.

### 3.5 Population genetic analyses based on 55 AISNPs

We first calculated the pairwise Cavalli-Sforza genetic distance between 166 populations based on the 55 Kidd AISNPs. As shown in **Fig. S18** and **Table S16**, genetic clusters were associated with language classification or geographical location. The smallest genetic distance was identified between Haikou Han and Meizhou Hakka (0.0049), followed by distances from Guangdong Han (0.0061), Guangzhou Han (00065) and Kinh (0.0082). Qiongzhong Hlai harbored the smallest genetic distance from Li (0.0052), followed by Dai (0.0106), Kinh (0.0162) and Ami (0.0203). Daozhen Gelao shared a close genetic relationship with southern Han (0.0039), followed by Yunnan Han (0.0047), Shaanxi Han (0.0084), and San Francisco Han (0.0086). **Fig. S19** shows the heatmap of pairwise genetic distance among the East Asians. The genetic distance between Xining Tibetans and other East Asians is large. Li and Qiongzhong Hlai clustered. Muli Tibetan, Dujiangyan Tibetan, Liangshan Yi and Ningxia Hui populations were clustered together, and Haikou Han and three southern Han groups were clustered. Beijing Han and Southern Han clustered with Daozhen Gelao.

We subsequently performed a principal component analysis based on the frequency distribution data of 55 AISNP genetic markers among 166 populations. As shown in **Fig. 4**, the first three components accounted for 87.03% of the total variance. The first component separated East Asians from other groups, and the second component separated Africans from other groups. A two-dimensional scatterplot constructed based on PC1 and PC2 (**Fig. 4A**) revealed that Africans, Europeans, and Central and South Asians were clustered to form a genetic cline, starting from the African population in the negative Y-axis direction and ending in the European group in the positive Y-axis direction. Eurasian populations and American populations clustered to constitute the Eurasian genetic cline, starting from the European populations in the negative direction of the X-axis and ending with the East Asians in the positive direction of the X-axis. Xining Tibetan and Kumul Uyghur populations were clustered with Central Asians. Ningxia Hui, Liangshan Yi, Muli Tibetans and Dujiangyan Tibetans were clustered with the Tibetan-Burman-speaking groups, and Haikou Han, Qiongzhong Hlai and Daozhen Gelao were clustered with southern Chinese populations. The third component separated the American populations from other populations. In the two-dimensional scatterplot constructed based on the first and third components (**Fig. 4B**), American populations were clustered in the upper right corner, and East Asians were clustered in the lower-right corner. Newly studied populations were grouped into three main clusters. In the two-dimensional scatterplot based on the second and third components (**Fig. 4C**), Americans and Africans were clustered and separated from other populations. Among 56 East Asians (**Fig. 4D**), the top ten principal components revealed 89.74% of the variance. In the two-dimensional scatterplot of East Asians, groups from the same language were generally clustered together. Austronesian and Tai-Kadai speakers were clustered in the lower-left corner to with the Qiongzhong Hlai. Haikou Han, Daozhen Gelao and Chinese-related groups gathered in the upper left corner. Ningxia Hui, Muli Tibetan, Dujiangyan Tibetan and Liangshan Yi populations were clustered with Tibetan-Burmese groups. Xining Tibetan and Kumul Uyghur populations were scattered and located on the far right.

**Fig. 4.**
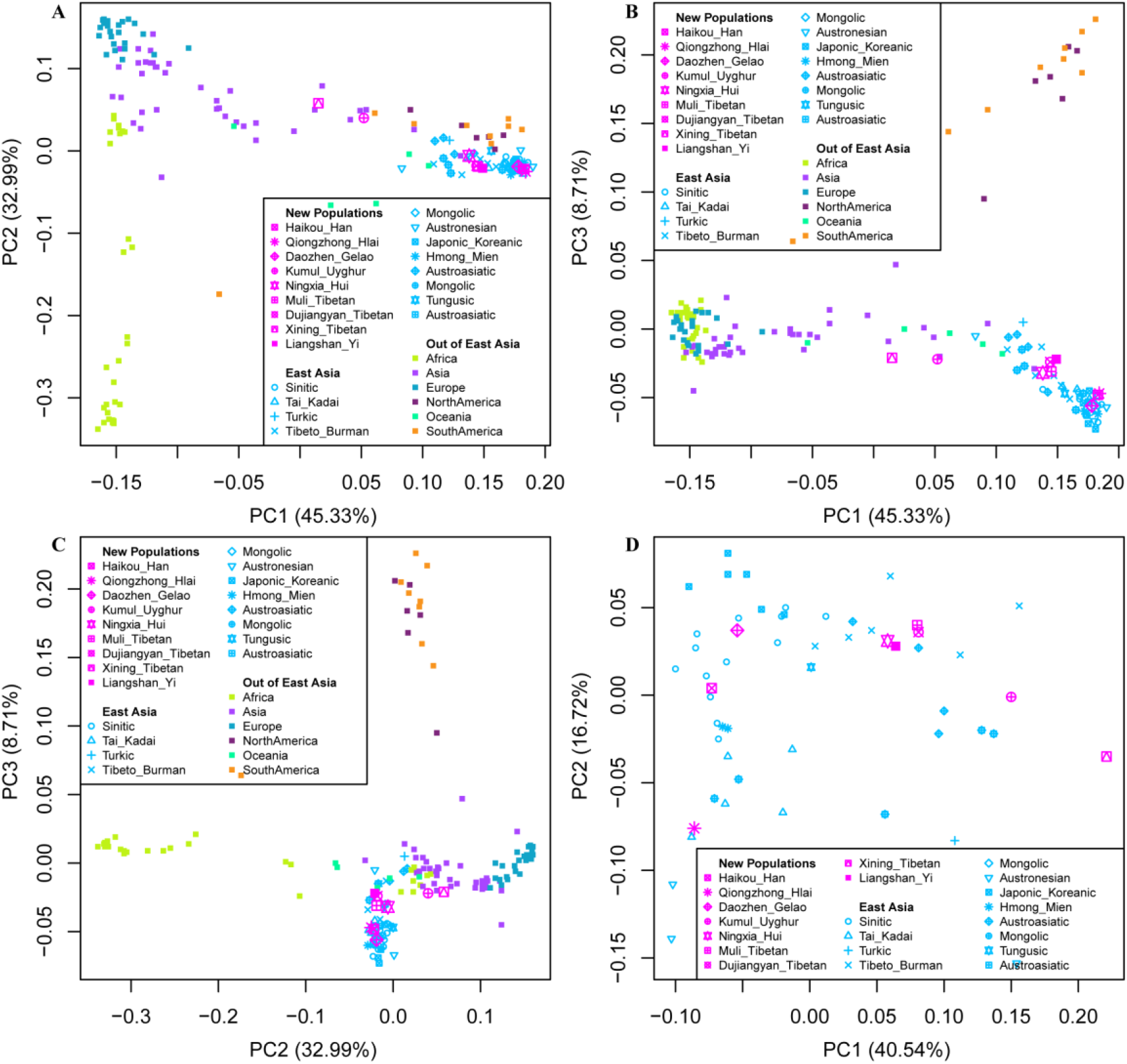
Patterns of genetic relationships among 166 populations worldwide. (**A∼C**). Principal component analysis among 166 populations worldwide based on allele frequency distributions of Kidd 55-SNP AIMset. (**D**). PCA showing genetic affinity within East Asians. East Asian subset PCA was conducted separately.

Finally, we reconstructed the neighbor-joining-based phylogenetic tree of the 166 included global populations (**Fig. 5**). Four large cluster branches were identified: the African branch, western Eurasian branch, American branch and East Asian branch. We observed that some populations from the Arabian Peninsula were clustered into the African branch, while Europeans and most of the Central and South Asians were clustered and constituted the western Eurasian branch. Clusters of Americans constituted the American branch. East Asian branches could be divided into two subbranches. One consisted of Mongolic, Tungusic, and Turkic speakers in the Tans-Eurasian language family, Tibetan-Burmese and Oceanian groups. Other East Asian groups, including Sinitic, Hmong-Mien, Tai-Kadai, Austronesian and Austroasiatic speakers, were clustered and formed the other subbranch. Qiongzhong Hlai first clustered with the Austronesian-speaking Ami, Atayal and Tai-Kadai-speaking Li and then with the Dai. The Haikou Han population first clustered with the Guangxi Miao, Guizhou Miao and Guangxi Han and then clustered with the Meizhou Hakka and Southern Han. Daozhen Gelao clustered with Japanese groups.

**Fig. 5.**
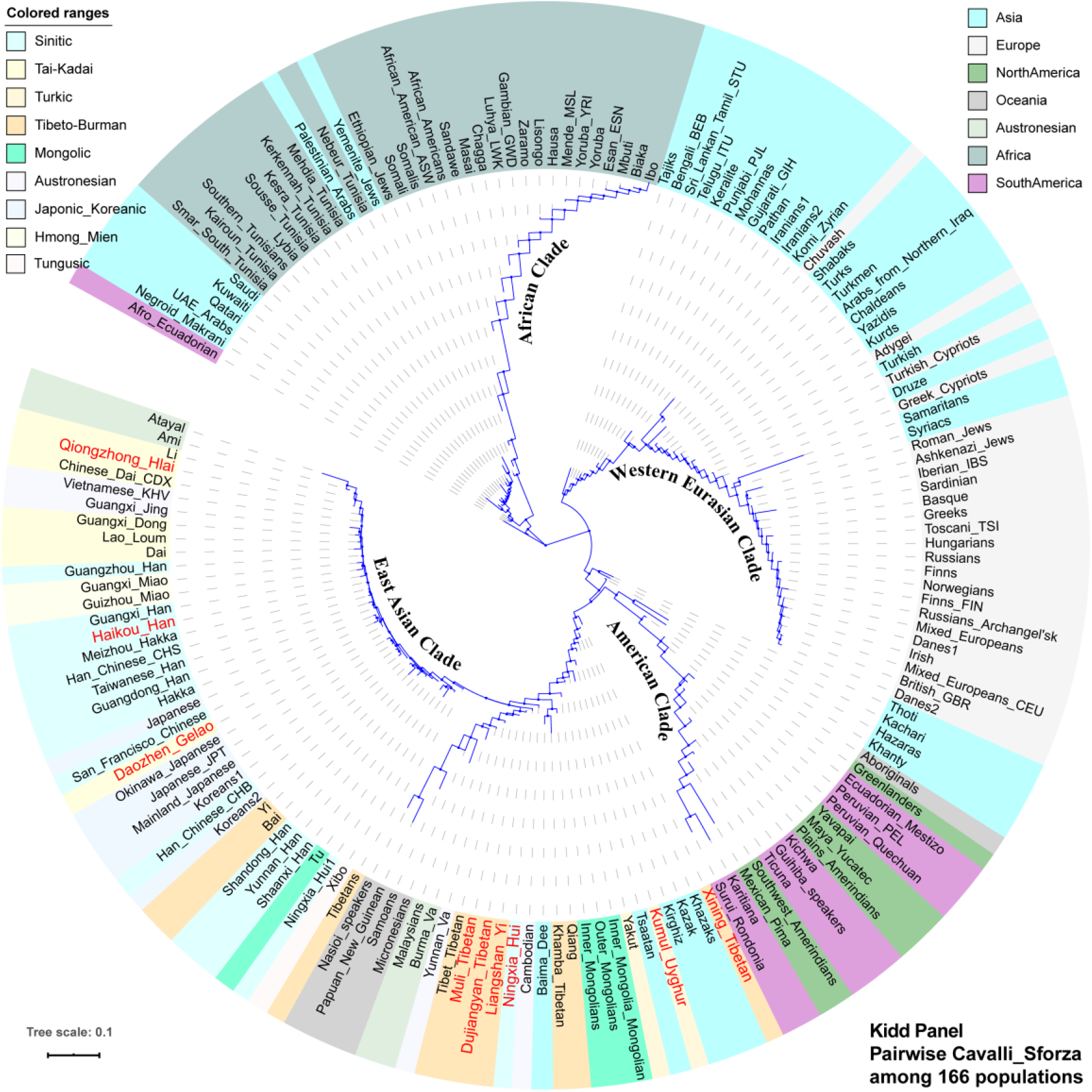
Neighbor-joining-based phylogenetic relationship among 166 populations worldwide based on Scali-Sforza genetic distances. All included populations are color-coded by geographical divisions or linguistic affiliation.

### 3.6 Genetic structure analyses based on 123 Seldin AISNPs

Subsequently, we merged our data with the previously published Seldin dataset and finally obtained a new merged data set consisting of 123 AISNPs from 140 populations worldwide. We first estimated pairwise Cavalli-Sforza genetic distances. As shown in **Fig. S20** and **Table S17**, the pairwise genetic distance between Haikou Han and Daozhen Gelao was the smallest (0.0067), followed by distances from Hakkas (0.0085), San Francisco Chinese (0.0090) and Chinese Han (0.0096). The minimum pairing genetic distance between Qiongzhong Hlai and Lao Loum was 0.0134, followed by distances from Haikou Han (0.0137), Daozhen Gelao (0.0142) and Hakkas (0.0151). Daozhen Gelao showed a close genetic affinity with the San Francisco Chinese (0.0051), followed by Chinese Han (0.0058) and Hakkas (0.0061). Daozhen Gelao and Haikou Han clustered together and showed a close genetic affinity with Lao Loum, and Qiongzhong Hlai clustered close to Filipinos in the heatmap of the worldwide populations, which was consistent with the common origin of Tai-Kadai- and Austronesian-speaking populations revealed by genome-wide chip data [25]. Among 55 East Asians (**Fig. S21**), a clearer close genetic relationship could be identified between Tai-Kadai speakers (Li, Qiongzhong Hlai) and Austronesian-speaking Filipinos.

A total of 89.39% of the variance was extracted from the 140 worldwide populations via the top ten PCs. As shown in **Fig. S22**, PC1 separated the American populations from other reference populations, and PC2 separated Africans from other populations, differing from the results based on the Kidd dataset. In the PC1-PC2 plot, Africans were clustered and constituted the African genetic cline, Europeans were clustered in the lower-left corner to form the European cluster, East Asians were clustered and formed the East Asian cluster, and Americans were clustered and formed an American cluster. South and Central Asians were clustered and located between the European cluster and the East Asian cluster. Ningxia Hui and Kumul Uyghur populations were clustered with South and Central Asians. Gelao, Hlai and Haikou Han were located close to East Asians. In the PC1-PC3 plot, Americans were separated and clustered in the upper right corner, and the other groups were clustered into genetically gradual groups that began with Europeans and Africans and ended in East Asians. In the PC2-PC3 plot, Africans were separated and clustered on the right side, and Americans were separated and clustered in the upper area. The cluster in the lower-left corner formed the Eurasian genetic cline, starting from Europeans, extending to Central and South Asians, and ending in East Asians. Focusing on 44 populations in East Asia, the top ten components could extract 68.73% of genetic variation. In the two-dimensional East Asian scatterplot, the Qiongzhong Hlai people clustered with the Vietnamese, while the Haikou Han people were grouped with the Hakkas and Han peoples. Daozhen Gelao, Liangshan Yi and HGDP Yi were clustered, while Muli Tibetan, Dujiangyan Tibetan and Tujia populations were gathered.

Phylogenetic relationships reconstructed based on Seldin’s dataset (**Fig. S23**) similarly showed four major clustered branches: the Eastern Eurasian clade, American clade, African clade and western Eurasian clade. The African clade was mainly formed by populations from Sub-Saharan Africans. American populations were clustered to form the American clade. The western Eurasian clade was mainly formed by clusters of Europeans, some North Africans, and Central Asian and Arabian Peninsula groups. The eastern Eurasian clade was clustered by East Asians and some Central Asians. Three groups in the present study were clustered with the East Asian terminal branch in the Eastern Eurasian clade. Qiongzhong Hlai and Ami, Atayal, Li and Filipinos first clustered together, and then these groups clustered with Haikou Han. Daozhen Gelao and Hakkas clustered first.

## 4 Discussion

### 4.1 Evaluating the efficacy of forensic ancestry inference

The individual ancestral inference efficiency of the Precision ID Ancestry Panel has been validated in various populations worldwide [3, 4, 12-15]. However, the related inference efficacy in southern Chinese populations has not been comprehensively evaluated. A total of 165 AISNPs in 691 samples from 9 populations in China were sequenced, 361 individuals from three southern Chinese populations were first reported here, and the efficiency of ancestral inference for 165 AISNPs was comprehensively evaluated.

For the ancestry inference of the East Asians, based on the built-in analysis platform (HID_SNP Genotyper plug-in) and model-based ADMIXTURE, admixture proportions of the tested individuals were inferred, and the specific admixture ancestral sources of the individuals were predicted. The built-in ancestral composition ratio is inferred based on the genetic variation information among populations from Africa, Europe, Southwest Asia, South Asia, East Asia, Oceania and America. Because the reference groups also included genetic reference data from southern East Asia, such as the southern Han Chinese, Hakkas, and Ho Chi Minh Kinh in Vietnam, three studied populations (Daozhen Gelao, Haikou Han and Qiongzhong Hlai) can be more accurately assigned to their origins in East Asia using the built-in ancestry assignment algorithm. When inferring the individual ancestral composition of Tibetan-Burmese groups, although it can be more accurately judged that their ancestral components were mainly derived from East Asians, the admixture proportion of South Asian and American related ancestry has gradually increased. For the ancestral assessment of the northwestern groups, such as the European-Asian admixed Uyghurs, the sources of their individual ancestral components are more complicated. When inferring the source of an individual’s ancestry based on the likelihood ratio, an investigated individual may be mismatched to other ethnic groups in the vicinity or to the same ethnic group from different geographical locations.

Forensic biogeographic ancestry inference is highly dependent on four major factors: (I) the understanding of the fine-scale genetic structure map of culturally/geographically/ethnolinguistically diverse global/regional populations, which was the basis for designing the AISNP panel and establishing a reference database; (II) the development and validation of an ancestry inference panel with global resolution and local regional discrimination; (III) the accomplishment of a global high-coverage population reference database; and (IV) the development of forensic-specific statistical methods. Thus, to improve the efficiency of ancestry inference among East Asians, we provide three recommendations. First, we should establish a full-coverage reference population database of one newly developed or updated AIMset. It is necessary to comprehensively explore the characteristics of genetic diversity of different global populations (especially for groups with different genetic structures) so that the accuracy of biogeographical ancestry inference can be improved. Second, the flaws of ancestral inference-related statistical methods must be overcome. At present, many comprehensive ancestry inference methods are being optimized and validated, such as allele-shared-based *f*-statistics, local ancestral inference, and IBD matrix ancestral inference, but these methods were initially dependent on the high density of genetic markers and mainly used for molecular anthropologic studies [38]. Third, we should develop and validate the East Asian-specific AIMsets panel. This study and previous genetic surveys have found that East Asians have a complex demographic history and differentiated population structure [19, 22-24, 41]. Thus, updated region-specific AISNP panels need to be developed.

### 4.2 Genetic structure and population relationships

In the exploration of population genetic relationships among 112 global populations based on the original genotype data of the Precision ID Ancestry Panel with the exception of rs10954737, PCA could divide the 112 included populations into five intercontinental groups: Africans, Europeans, East Asians and Native Americans and South Asians, consistent with the clustering pattern inferred from the Human Genome Diversity Project [42]. In the ADMIXTURE analysis results, we observed the smallest cross-validation error value when K = 5 among worldwide populations and Eurasian groups. The ancestral component analysis in the 1000 Genomes Project set was able to separate further the South Asian and American clusters as the predefined ancestral sources increased. At the same time, we also noticed that the Medellin Colombian and Puerto Rican populations in the Americas showed obvious admixed genetic backgrounds consisting of African ancestry, European ancestry, Native American ancestry and South Asian ancestry. Model-based ADMIXTURE analyses among Eurasians and East Asians showed that northwestern Chinese populations, including the Kumul Uyghur, Xining Tibetan and Xinjiang Kazakh peoples, were modeled as an admixture of East Asian-related ancestry, European-related ancestry and South Asian-related ancestry. These observed patterns of genetic admixture were consistent with the results of an earlier population genomic study focused on Xinjiang Uyghurs. Xu et al [43] genotyped 951 Xinjiang Uyghur individuals from 14 groups using whole-genome chip technology. Their results showed that the Uyghur people was formed by mixing the two ancestral populations related to eastern and western Eurasians during the Bronze Age. Western Eurasian ancestry consisted of a mixture of 25-37% European-related ancestral components and 12-20% South Asian-related ancestral components. Eastern Eurasian ancestry was formed by mixing 15–17% of the Siberian-related ancestral component and 29-47% of the East Asian-related ancestral component. Our study did not simulate the Siberian ancestral component, because the original genotype data lacked northern Siberian population data. We also found that northwestern Tibetans in Qinghai Province tended to cluster with the aforementioned Uyghur and Kazakh peoples, which was consistent with the population stratification among culturally diverse Tibetans and western Eurasian affinity of Ando Tibetans in the Ganging region revealed via ancient/modern genome-wide data and Y chromosome genetic variations [22, 44].

Additionally, we used three TreeMix-based phylogenies to explore the phylogenetic split and potential gene flow events and to verify the patterns of population genetic relationships inferred from the PCA and ADMIXTURE. Among worldwide populations, the Medellin Colombian and Puerto Rican clusters were located between South Asian and European groups. Lima Peruvians and Los Angeles Mexican Americans shared a closer phylogenetic relationship with East Asians. Among Eurasian or East Asians, we observed a strong association between genetic affinity and linguistic affiliation. The newly studied Chinese populations can be basically divided into three subgroups: the northwestern cluster included Mongolic, Tungusic and Turkic speakers; the Tibeto-Burman cluster and southern Chinese cluster consisted of Sinitic, Austronesian, Tai-Kadai and Hmong-Mien speakers, which is also consistent with the geographic classifications (South, West and Northwest China). The three studied southern populations (Han, Gelao and Hlai) were included in the southern Chinese cluster. These phylogenetic relationships can also be confirmed via two N-J-based phylogenetic trees using Cavalli-Sforza genetic distance matrixes with Kidd’s dataset and Seldin’s dataset. Focusing on admixture events and gene flow, we found that approximately 40% of Finnish-related ancestry was mixed into Kumul Uyghurs, and 36% of European-related ancestral components were mixed into the Xinjiang Kazakh gene pool. PCA and pairwise genetic distance results based on the Kidd AISNP sets from 166 populations revealed an African genetic cline and a west-east Eurasian cline. However, the American population was first to be clearly separated among 140 populations based on Seldin’s AISNP sets. This result was consistent with the initial development purpose of Seldin’s inference system, focused on the distinction between Americans and others. The finer-scale genetic admixture history of East Asians should be further explored via high-density variants or whole-genome sequencing data in the next step. At the same time, the complex genetic admixture history and population substructure of East Asians observed in our study further reminded East Asian forensic scientists that the fine-scale genetic structure of East Asians is not fully understood at present; thus, comprehensive dissection of the East Asian population substructure and development of an East Asian-specific ancestry inference system combining the distinguishing resolution of the global population and high discrimination within East Asian subpopulations is an urgent goal.

## 5. Conclusion

Our study provided the first batch of forensic reference population data from 316 individuals from southern Chinese Sinitic- or Tai-Kadai-speaking populations. The estimated forensic parameters showed that the Precision ID Ancestry Panel was developed for the purpose of biogeographical ancestry inference. Its forensic performance for personal identification is comparable to the power of the currently widely used Huaxia Platinum STR system. Based on cluster comparisons from three different datasets and ancestral source inference based on the HID_SNP Genotyper plugin, ADMIXTURE and PCA, we found that this AISNP system had a strong intercontinental resolution, but greater instability and mismatch rate remained in the intracontinental differentiation of subpopulations. Preliminary population analysis results showed three different substructure groups in the newly studied Chinese populations: a Southern Chinese cluster, a Tibetan-Burmese cluster, and a northwestern cluster, which showed a western Eurasian admixture. Population admixture and split models in TreeMix analyses revealed that Kumul Uygur and Xinjiang Kazakh populations were admixtures of approximately 40% European-related ancestry and 60% East Asian-related ancestry. In addition, comprehensive population genetic analyses further demonstrated that the Tai-Kadai-speaking Hlai people harbored a homogeneous genetic structure and possessed genetic affinity with Austronesian populations; thus, their ancestry can be used as an ancestral source proxy in subsequent population genetic history reconstruction of southern Chinese and as a representative for forensic database establishment of East Asians.

In summary, this study found that the AISNP system has the potential for personal identification, but its forensic ancestry inference capacity needs to be further optimized for the composition of included East Asian-specific AIMsets and the establishment of a more comprehensive reference comparison database. In addition, detailed exploration of the genetic structure of East Asians and the development of a region-focused ancestry inference system that combines the distinguishing resolution of global populations and high discrimination within East Asians are the two key issues that need to be resolved for accurate ancestry inference in East Asians.

## Conflict of interest

The authors declare that they have no conflict of interest.

## Acknowledgements

We would like to thank the volunteers who contributed samples for this study. This study was supported by grants from the National Natural Science Foundation of China (81501635) and the Fundamental Research Funds for the Central Universities (YJ201651). The funders had no role in study design, data collection and analysis, decision to publish, or preparation of the manuscript.

## Legends of Supplementary Figures

Table S1. Detailed sample information of studied populations and reference populations.

Table S2 Genotypic data of 165 AISNPs for three newly studied populations

Table S3. Allele frequencies and corresponding forensic parameters in the Daozhen Gelao population.

Table S4. Allele frequencies and corresponding forensic parameters in the Haikou Han population.

Table S5. Allele frequencies and corresponding forensic parameters in the Qiongzhong Hlai population.

Table S6. Pairwise F_st_ genetic distances among 112 worldwide populations based on the 164 AISNPs.

Table S7. Ancestry proportions estimated via ADMIXTURE analysis of 6933 individuals from 112 populations with the predefined ancestral sources ranging from two to twenty.

Table S8. Ancestry proportions estimated via ADMIXTURE analysis of 4506 individuals from 67 populations with the predefined ancestral sources ranging from two to twenty.

Table S9. Ancestry proportions estimated via ADMIXTURE analysis of 1993 individuals from 29 populations with the predefined ancestral sources ranging from two to twenty.

Table S10. Admixture signals for Daozhen Gelao via *f*_*3*_*(Source1, Source2; Daozhen Gelao)*.

Table S11. Admixture signals for Haikou Han via *f*_*3*_*(Source1, Source2; Haikou Han)*.

Table S12. Admixture signals for Qiongzhong Hlai via *f*_*3*_*(Source1, Source2; Qiongzhong Hlai)*.

Table S13. Results of genetic affinity inferred from symmetrical-*f*_*4*_-statistics in the form *f*_*4*_*(Reference population1, reference population2; studied populations, Mbuti)*.

Table S14. Results of genetic affinity inferred from symmetrical-*f*_*4*_-statistics in the form *f*_*4*_*(Reference population1, studied populations; reference population2, Mbuti)*.

Table S15. Results of shared derived alleles inferred from symmetrical-*f*_*4*_-statistics in the form *f*_*4*_*(studied population1, studied population2; reference population, Mbuti)*.

Table S16. Pairwise Cavalli-Sforza genetic distances among 166 populations worldwide based on the Kidd 55-SNP AIMsets

Table S17. Pairwise Cavalli-Sforza genetic distances among 140 populations worldwide based on the Seldin 123-SNP AIMsets.

## Notes

### Competing Interest Statement

The authors have declared no competing interest.

